# Synergistic effects of dietary taurine and carbohydrates supplementation on skeleton muscle of juvenile turbot *Scophthalmus maximus*

**DOI:** 10.1101/2023.05.10.540249

**Authors:** Hasi Hays, Wenbing Zhang, Kangsen Mai

**Author notes:** **Correspondence:** (H.H.), (W.Z.).

## Abstract

The present study aims to investigate the effects of dietary taurine and carbohydrate levels on the skeleton muscle growth of turbot. Muscle samples of turbot were collected after 70 days of feeding trial by treatment groups of 0% (C), 0.4% (L1), 1.2% (L2) taurine with 15% dietary carbohydrate level, and 0.4% (H1), 1.2% (H2) taurine with 21% dietary carbohydrate level. Results showed that L2 and H2 treatment has given significantly higher hyperplasia with significantly high muscle fiber frequencies and muscle fiber density than that in the other groups. Hyperplastic muscle fiber generation was significantly stimulated by the high carbohydrate level (21%). Muscle density was not dependent on the level of carbohydrates. Aspartate, Threonine, Serine, Glutamine, Leucine, Phenylalanine, Isoleucine, Lysine, Histidine, and Arginine were significantly high in the H1 group than that in other all groups. H2 treatment was given a significantly higher amount of total collagen content than the other groups by increasing alkaline-soluble, alkaline-insoluble hydroxyproline, and total hydroxyproline levels. Hardness has significantly increased in all the treatment groups than that in the control group. And also, muscle hardness was significantly increased by the dietary carbohydrate levels. Intestine amylase, lipase, and trypsin enzyme activities were significantly increased in all the treatment groups than that in the control. Amylase and lipase activities were significantly highest in the H2 group. Taurine 1.2% with carbohydrates 21% treatment group (H2) was given significantly higher levels of cellular level muscle growth with more collagen in the skeletal muscle of Turbot.

## 1 Introduction

Aquaculture is the main protein source in human nutrition. Aquaculture production has increased in past decades worldwide to supply world demand. Hence, fishmeal has used as the main protein in the fish diet. But due to the high cost and less availability of fish meals, the industry has looked forward to alternative protein sources (FAO, 2018). Taurine (2-aminoethanesulfonic acid) is typically found in relatively high concentrations in fish meals and animal by-products but is almost non-existent in plant meals. Even when all essential amino acid requirements are met in plant-based diets for carnivorous fish, growth still is often reduced when compared to fish meal-based diets (Gaylord et al., 2006). Taurine is abundant in animal tissues and freely available amino acid. Taurine is not considered to be an essential amino acid in fish because it can be synthesized in the liver. It is a conditionally essential amino acid in fish diet. The optimum taurine requirement mainly depends according to the fish species (Kim et al., 2005; Sampath et al., 2020a; Sundararajan et al., 2014). In the mammalian system, taurine is synthesized through many enzymatic reactions, but the enzyme L-cysteine-sulphinate decarboxylase appears to be rate-limiting. But in fish taurine is synthesized from L-cysteine after the process of oxidative enzymatic action in the biosynthesis process in the liver (Jacobsen and Smith, 1968; Yokoyama et al., 1997; Liu et al., 2017). The activity of this enzyme varies in fish depending on species and size. For example, in the yellowtail, as well as in bluefin and skipjack tunas, L-cysteinesulphinate decarboxylase activity is not present, whereas in Japanese flounder it expresses only low activity (Yokoyama et al., 2001).

Fish raised in seawater may have a greater demand for dietary taurine than fish held in freshwater and the ability of the fish to convert cysteine to taurine may be based on their environmental salinity requirements and other environmental factors (Gaylord et al., 2006). Turbot (*Scophthalmus maximus*) is a carnivorous marine fish that is greatly affected by taurine supplementation by optimizing the growth performance. Dietary taurine additions improve weight gain and feed efficiency in olive flounder (Sampath et al., 2020a; Park et al., 2002; Kim et al., 2005), as well as in rainbow trout (Gaylord et al., 2006). Taurine improves the growth performance of several fish species including turbot (*Scophthalmus maximus*) (Wei et al., 2018; Liu et al., 2017, Zhang et al., 2019), barramundi (*Lates calcarifer*) (Poppi et al., 2018), black carp (*Mylopharyngodon piceus*) (Zhang et al., 2018), red sea bream (*Pagrus major*) (Takagi et al., 2010), California yellowtail (*Seriola lalandi*) (Salze et al., 2018), olive flounder (*Paralichthys olivaceus*) (Kim et al., 2017), and yellowtail (*Seriola quinqueradiata*) (Nguyen et al., 2015). Therefore, taurine supplementation may be required for plant-based diets and indeed, and also in fish meal diets for the optimum growth performances of the fish. Taurine was reduced the enzyme activity of glucogenesis, amino acid, and lipid peroxidation in the liver (López et al., 2015). And also, digestive enzyme activity has key roles in nutrition digestion in the intestine. Dietary taurine has significantly increased the amylase and lipase activity in the intestine of black carp (*Mylopharyngodon piceus*). But amylase and lipase activity levels were significantly reduced without taurine supplementation. So, taurine has key roles in nutrition digestion that is important in optimum fish muscle growth (Zhang et al., 2018).

The combined effects of taurine and carbohydrates supplementation have improved the growth performances, enzymatic activities, muscle taurine, blood glucose levels, and oxidation activities of the turbot. Intestinal amylase, mRNA levels of liver phosphofructokinase, glucokinase, glucose-6-phosphate dehydrogenase (G6PD), pyruvate kinase, glycogen synthase (GS) has significantly improved with 1.2% taurine supplementation with the 21% dietary carbohydrates level of turbot (*Scophthalmus maximus L*.). At the same time, mRNA levels of liver cytosolic phosphoenolpyruvate carboxykinase (cPEPCK) have significantly reduced by dietary taurine levels (Zhang et al., 2019). But the combined effects of taurine and carbohydrates for muscle growth parameters and histological effects have been less investigated. So, the current study has focused on the cellular level muscle growth of turbot (*Scophthalmus maximus*) fed diets with taurine and carbohydrates supplementation by measuring the cellular level muscle fiber development via hyperplasic and hypertrophic histochemical analysis and nutrition digestion via digestive enzyme activity levels.

## 2 Materials and methods

### 2.1 Experimental diet

The present study was conducted with specially prepared formulated feed for turbot dietary requirements. The diet was formulated with two levels of carbohydrates and two levels of taurine levels. Furthermore, the experimental diets were formulated as 15% of carbohydrates with supplementing 0.4% (L1), 1.2% (L2) taurine and 21% of carbohydrates with supplementing 0.4% (H1), 1.2% (H2) taurine. The control diet (C) was formulated with 15% carbohydrates and no supplementation of taurine.

The basal diet was contained 50% crude protein, 15% carbohydrates and 11% crude lipid for Turbot. The formulated taurine powder was contained 99.2% purity. (Yongan Pharmaceutical Company Limited, Qianjiang, China. Batch No: STP14042149). All the dry ingredients were sieve by 500 μm mesh before mixing.

All the diets were extruded into two sizes 1.2 mm and 2.2 mm. Finished feeds were stored at -20 °C until the start of the feed trial.

### 2.2 Feeding trial

Turbot (Average body weight 3.66 ± 0.02 g) was purchased from the commercial breeding center at Weihai, Shandong, China. Juvenile turbot was fed with diet C for 14 days in a laboratory condition to acclimatize to the same experimental conditions. Feeding trial was done in 15 fiberglass tanks with 300 L water capacity with the indoor flow-through system at Yellow Sea Fisheries Institute, Yantai, China. Each tank was included 40 healthy fish. And also, each treatment has consisted of triplicate tanks. Tanks were fully aerated with oxygen and followed natural light and dark routine methods. Feeding was done twice a day and mortality, dissolved oxygen (DO), pH, salinity, and water temperature was recorded daily. Average water quality parameters of dissolved oxygen (DO), pH, salinity, and the water temperature were noted 7.0 mg l^-1^, 7.5-8.0, 30-33, and 15-20 °C respectively. Tanks were cleaned every day to maintain sanitary condition throughout the experiment. Tanks were covered with a wire mesh to retain fish inside the tank. Fish were randomly selected and taken body weight 10 fish per tank weekly basis to ensure the feeding intake. All the experiment guidelines were strictly followed with the Guide for the Use of Experimental Animals of the Ocean University of China.

### 2.3 Sample collection

Fasting regime was followed for 24 h before anesthetizing. Fish were anesthetized with MS-222 (Shanghai Reagent, China). Bodyweight was measured 15 fish per each tank and, skeleton muscle samples were collected from the dorsal region of 6 fish per each tank. One portion of each sample was stored in 4% paraformaldehyde for histochemistry analysis. Another portion was stored in liquid nitrogen and stored at - 80 °C for the analysis. And the other side of the fish fillet was used to measure Alkaline Soluble Hydroxyproline (Hyp), alkaline-insoluble Hyp, collagen content, muscle texture parameters, and amino acids.

### 2.4 Histochemistry

Skeleton muscle samples were sectioned into small square shape pieces. Each section was washed with graded sucrose series. Then each muscle section was embedded in hot paraffin and cool down into sloid paraffin cubes. Those paraffin cubes were cut (Leica-RM2235, Germany) into 8 μm thickness slices and deposited on microscope glass slides. Then the glass slides were coated by using Poly-L-lysine (by Sigma) and air-dried about 30 min. Each slide was stained with hematoxylin-eosin before the microscopic examination. Microscopic photographs of samples were taken for morphological analysis. Muscle fiber morphology was analyzed by using ImageJ (Version 1.52a). Individual Muscle fiber was manually marked and calculated the muscle fiber density (200 × 100 μm^2^) and muscle fiber diameter. More than 200 muscle fibers of each replicate were measured. In total, more than 3,000 muscle fibers (>200^*^3replicates^*^5treatments) were manually marked in ImageJ software to increase the accuracy of data. Muscle fiber diameter frequencies were calculated as a percentage of each muscle fiber diameter class (Asaduzzaman et al. 2017).

### 2.5 Collagen determination

Muscle samples were dissolved in alkaline solution to separate alkaline soluble and insoluble hydroxyproline. To achieve the above process, 1g of muscle samples were minced and followed by homogenization with 9 mL cold water 1 min at 30,000 rpm. 0.2 M NaOH and 10 mL of ice-cold were added immediately. Then the wheel roller was used to mix the samples for 4 h (4 °C). The homogenate was separated and centrifuged 10,000 ×g for 30 min (4 °C) in ultracentrifuge (CR21GII, Hitachi, Japan). The remaining suspended liquid was analyzed for alkaline-soluble hydroxyproline (Hyp) content. Then, alkaline-insoluble hydroxyproline solid particles were re-suspended with 3 mL 6 M HCl. The re-suspended solution was transferred to a 5 ml ampoule bottle. The samples were hydrolyzed with distilled water up to 10 ml and kept for 20 h at 110 °C. The hydroxyproline was determined by the process described in Zhang et al. (2013). 2 mL of buffered chloramines T reagent (1.4 g chloramines T dissolved in 20 ml water and then diluted with 30 ml n-propanol and 50 ml acetate-citrate buffer (pH 6.5) were added to the hydrolyzed tissue sample and incubated for 20 min at room temperature. 2 ml of perchloric acid (18.9%) was added to the sample mixture and incubated for 5 min at room temperature. Then the 2 ml of 2 mL P-DMAB solution (10% w/v P-DMAB in n-propanol) was added to each sample and heated for 20 min at 60 °C. The mixture was cooled down to room temperature and absorbance was measured at 560 nm. The standard curve was used to determine the hydroxyproline content. Hyp content was multiplied by 8 to calculate the total collagen content of the turbot muscle (AOAC, 2000).

### 2.6 Amino acids analysis

Muscle samples were cut into small pieces and freeze-dried for 24 h. Each sample was chopped into a mash form by using a glass rod. 30 mg of each powdered muscle sample was hydrolyzed with 15 mL of 6 N HCl at 110 °C for 24 h. Filtered samples were put in the N_2_ (g) vaporization chamber to separate the nitrogenous compound in the sample. Then samples were hydrolyzed with 1 mL of 6 N HCN and mixed well to dissolve all the adhered compounds to the glass wall. The Automatic amino acid analyzer (L-8900, Hitachi, Japan) was used to measure the amino acids. Methionine and cysteine contents were measured separately by oxidizing the samples with performic acids at -10 °C for 3 h. then the samples were freeze-dried with deionized water. Then methionine and cysteine content were analyzed by methods used in Wei et al. (2016). The liver Cysteine level was analyzed the same above-mentioned method to compare the effects on taurine biosynthesis in the liver.

### 2.7 Muscle texture analysis

Turbot fillets were used to measure muscle textural properties. Three points were marked on the fillet above the lateral line, between the dorsal and tail. Textural parameters were measured by using a Texture analyzer (TMS-PRO, FTC, America) which was equipped with an 8 mm cylinder probe. Texture Profile Analysis (TPA) was analyzed by double compression of muscle. The test conditions were included two consecutive cycles of compression with a constant speed of 30 mm/min with the deformation 60% of the original length, and the initial force is 0.1 N. Adhesiveness, chewiness, cohesiveness, hardness and springiness were measured to get the muscle textural condition of the fish fillet (Ginés et al., 2004).

### 2.8 Digestive enzymes

The intestinal segments of three fish from each replicate group were sectioned and cleaned with the distilled water before enzyme activity analysis. Intestinal segments were stored in a 10 ml centrifuge tube under -20 °C until digestive enzyme analysis. For enzyme analysis, 0.5 g of each segment was measured and homogenized in 5 ml 0.9% cold saline solution. Samples were centrifuged 3,000 rpm for 10 min (4 °C), and supernatant were separated. All the supernatant was labeled and stored under -20 °C to measure the trypsin, lipase and amylase contents (Zhang et al., 2018).

### 2.9 Statistical analysis

Alkaline-soluble Hyp, Alkaline insoluble Hyp, collagen contents, Amino acid levels, muscle fiber diameter frequencies, and muscle fiber density data were analyzed by one-way of variance (ANOVA). Two-way ANOVA was used to identify the interaction between taurine and carbohydrates. The assumptions of normal distributions and homogeneity of variances were checked before analysis. The frequency distribution of fast skeletal muscle fibers in different class data was expressed in percentages that underwent square root transformation before statistical analysis. All ANOVA were tested at a 5% level (*P* < 0.05) followed by Tukey’s test in IBM SPSS Statistics software version 23.

## 3 Results

### 3.1.1 Body weight

The final body weights of turbot were 33.39 ± 0.68^a^ g, 34.54 ± 1.13^ab^ g, 38.02 ± 0.91^b^ g, 34.61 ± 0.39^ab^ g, and 35.51 ± 0.44^ab^ g in C, L1, L2, H1 and H2 groups, respectively with the levels of significance (Zhang et al., 2018). There was a significant interaction between taurine and carbohydrates in a two-way ANOVA analysis of body weight.

### 3.2 Effects on hyperplastic and hypertrophic growth of muscle fibers

Different taurine and carbohydrate groups gave clear variation in the muscle fibers. Histochemical sections are shown in Figure 1. Large numbers of hypertrophic fibers have shown in control, L1 and H1 groups. And also, hyperplastic fibers are largely shown in the L2 and H2. Dietary supplementation of taurine gave a large number of hyperplastic muscle fibers which are represented the new cell generation of juvenile turbot skeletal muscles with 1.2% taurine levels. And also, hypertrophic fibers which are represented the growth or the enlargement of the existing muscle fibers were high in control and 0.4% taurine diets fed turbot muscle. Histological qualitative data were confirmed by the muscle fiber diameter frequencies in different supplementary groups (Table 1). Treatment groups with 1.2% dietary taurine level (L2 and H2) gave significantly high hyperplastic (Class 10, diameter d ≤ 10 μm) than other groups and treatment groups with 0.4% dietary taurine level (L1 and H1) gave significantly high hypertrophic (Class 60 and 70 fibers, diameter d ≤ 50 μm) than L2 and H2 groups without depending the carbohydrates levels (*P* < 0.05). But there was significantly high hyperplastic muscle fiber in 21% carbohydrates levels than 15% levels (*P* < 0.05) (Figure 2). There was an interaction between carbohydrates and taurine for hyperplasia muscle fiber proliferation. So, hyperplastic muscle fiber generation was stimulated by the high carbohydrate levels. Muscle fiber density was significantly higher in 1.2% (L2 and H2) of the dietary taurine-supplemented group than that in the other groups (*P* < 0.05) (Figure 3). Thus, muscle fiber density was also altered by the interaction of high carbohydrate levels.

**Figure 1:**
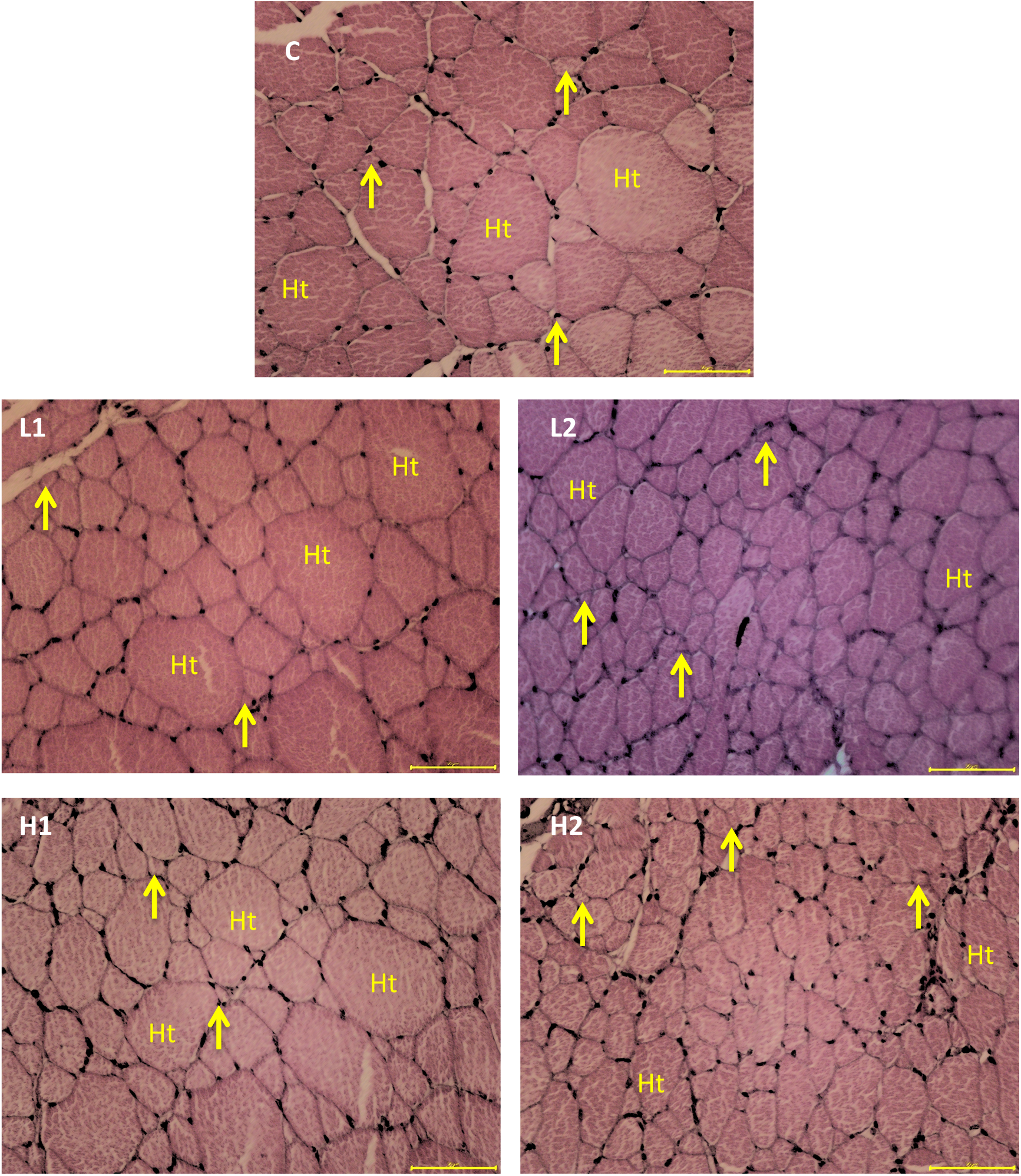
Histochemical sections of skeleton muscles of turbot after the 70-day feeding trial with treatment groups of 0% (C), 0.4% (L1), 1.2% (L2), 2.0% (L3) taurine with 15% dietary carbohydrate level, and 0.4% (H1), 1.2% (H2), 2.0% (H3) Taurine with 21% dietary carbohydrate level. Ht marks indicate the hypertrophic fibers and the arrowhead pointed to the hyperplastic fibers. Magnification is × 40 (Bar 50 μm).

**Table 1.** Mean level of muscle fiber diameter frequencies % effect by different levels of carbohydrates and taurine supplementations in turbot fed 70 days of feed trial.

|  |  | Muscle fiber diameter frequency classes |  |  |  |  |  |  |
| --- | --- | --- | --- | --- | --- | --- | --- | --- |
| CHO % | Taurine % | 10 | 20 | 30 | 40 | 50 | 60 | 70 |
| 15 | 0 | 6.358 | 36.67 | 26.07 | 19.867 | 4.471 | 2.928 | 3.637 |
| 15 | 0.4 | 3.511 | 33.858 | 28.652 | 13.479 | 16.425 | 0 | 4.076 |
| 15 | 1.2 | 14.762 | 31.524 | 27.667 | 17.143 | 5.714 | 0 | 3.19 |
| 21 | 0.4 | 12.625 | 34.075 | 29.663 | 11.513 | 3.838 | 2.762 | 5.525 |
| 21 | 1.2 | 15.439 | 41.668 | 25.37 | 11.119 | 2.649 | 2.649 | 1.105 |

**Figure 2:**
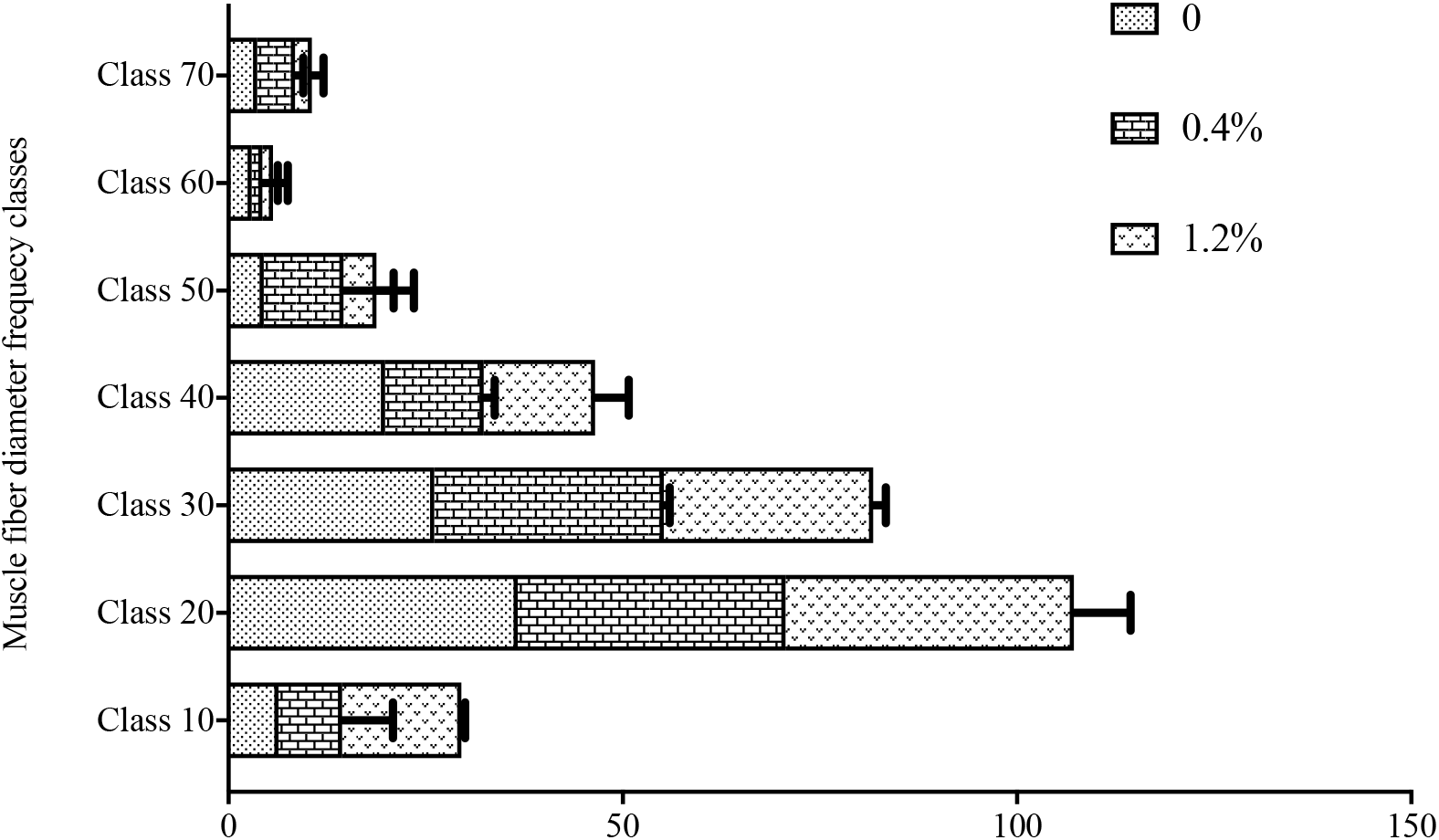
Frequency distribution of fast skeleton muscle fibers of turbot fed the diets supplemented with different levels of taurine for 70 days. Diameter classes (d, μm): class 10 = d ≤ 10, class 20 = 10 < d ≤ 20, class 30 = 20 < d ≤ 30, class 40 = 30 < d ≤ 40, class 50 = 40 < d ≤ 50, class 60 = 50 < d ≤ 60 and class 70 = d >60. The percentage values (One-way analysis of ANOVA) represent the bars and the top of the bar indicates the different superscript letters which is indicate the significant difference among the groups at 0.05 (*P* < 0.05). If the effects were significant, ANOVA was followed by the Tukey test. Error bars represent the deviation of 21% carbohydrate treatment groups from the 15% carbohydrate treatment groups that are represented by horizontal bars.

**Figure 3:**
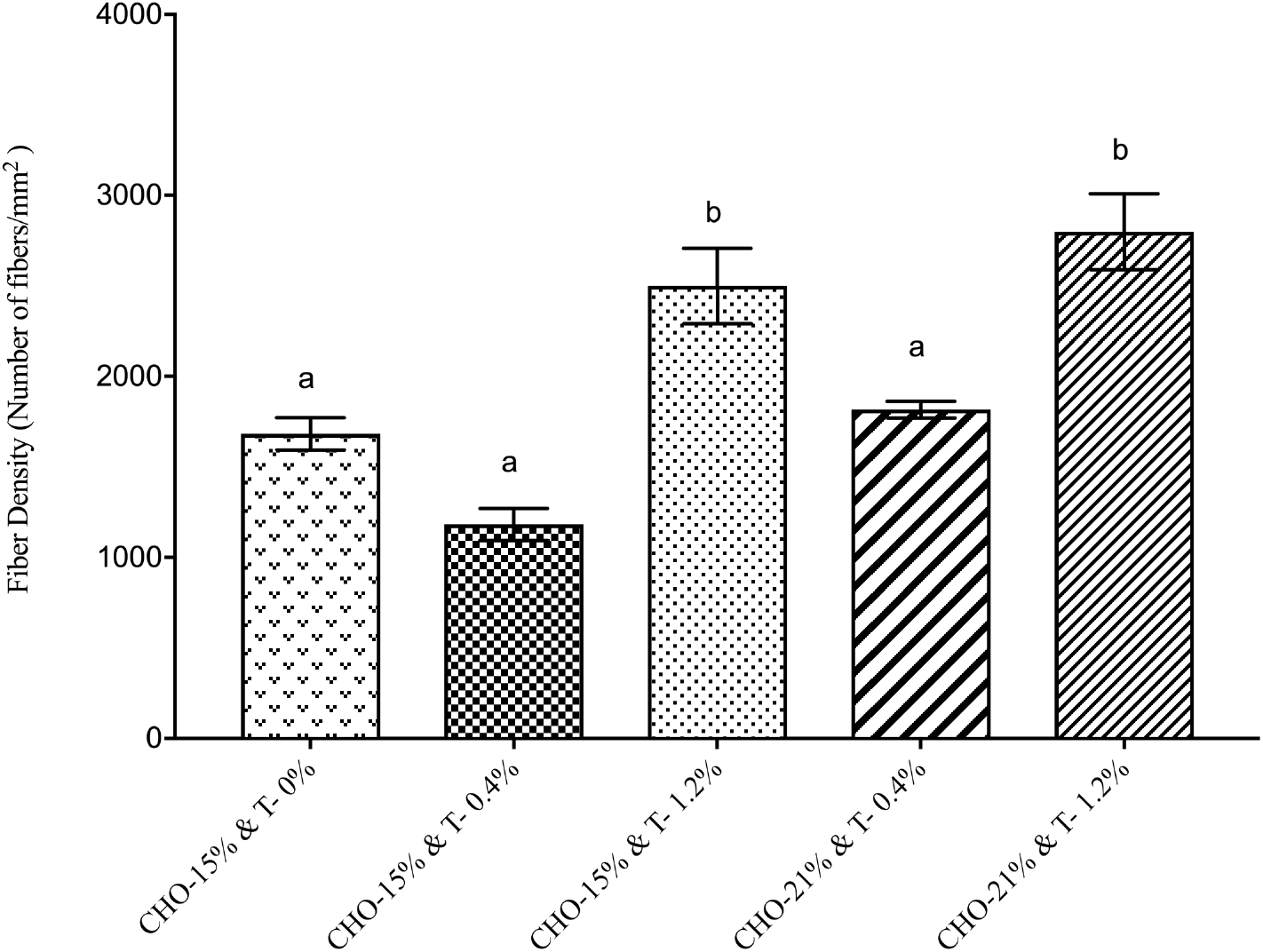
The density of fast skeletal muscle fibers of turbot fed the diets supplemented with different levels of taurine for 70 days. Values are means (±SE) of three replicates (each replicate consists of values from three samples) in each treatment. The mean values followed by the different superscript letters in each sampling data indicate a significant difference at 0.05 (*P* < 0.05). If the effects were significant, ANOVA was followed by the Tukey test.

### 3.3 Collagen contents

Alkaline-soluble hydroxyproline content, alkaline-insoluble hydroxyproline content, total hydroxyproline content, and total collagen are shown in Table 2. Alkaline-soluble hydroxyproline content in muscle has significantly increased with the taurine content as well as carbohydrates contents than that in the controls (*P* < 0.05). The highest hydroxyproline contents are shown in H2 (1.2% taurine and 20% carbohydrates) treated group. And also, alkaline-insoluble hydroxyproline content in muscle was significantly increased in the groups with 1.2% taurine and 20% carbohydrates (H2) levels (*P* < 0.05). The group with 1.2% of dietary taurine supplementation with 21% carbohydrates group had the significantly highest values of alkaline-soluble hydroxyproline content (0.32 mg/g), Alkaline-insoluble hydroxyproline content (0.29 mg/g), total hydroxyproline (0.61 mg/g) and the total collagen (4.88 mg/g) among all the groups (*P* < 0.05) (Figure 4). In two-way ANOVA analysis, there was an interaction between carbohydrates and taurine for the total collagen content of the muscle.

**Table 2.** Hydroxyproline (Hyp) and collagen content in the muscle of turbot after the 70-day feeding trial.

| Description<br>(mg /g wet<br>tissue) | C | L1 | L2 | H1 | H2 |
| --- | --- | --- | --- | --- | --- |
| Alkaline-soluble Hyp | 0.10 ± 0.008<sup>a</sup> | 0.17 ± 0.008<sup>b</sup> | 0.21 ± 0.017<sup>bc</sup> | 0.23 ± 0.008<sup>c</sup> | 0.32 ± 0.012<sup>c</sup> |
| Alkaline-insoluble Hyp | 0.23 ± 0.009<sup>a</sup> | 0.22 ± 0.018<sup>a</sup> | 0.27 ± 0.009<sup>c</sup> | 0.24 ± 0.012<sup>bc</sup> | 0.29 ± 0.012<sup>c</sup> |
| Total Hyp | 0.33 ± 0.015<sup>a</sup> | 0.39 ± 0.023<sup>b</sup> | 0.48 ± 0.009<sup>c</sup> | 0.47 ± 0.003<sup>c</sup> | 0.61 ± 0.003<sup>d</sup> |
| Total collagen | 2.62 ± 0.122<sup>a</sup> | 3.12 ± 0.187<sup>b</sup> | 3.84 ± 0.071<sup>c</sup> | 3.76 ± 0.027<sup>c</sup> | 3.88 ± 0.027<sup>d</sup> |

All data were expressed as mean ± SE. Mean values within the same row with different superscripts are significantly different ( $P < 0.05$ ; Tukey's test).

**Figure 4:**
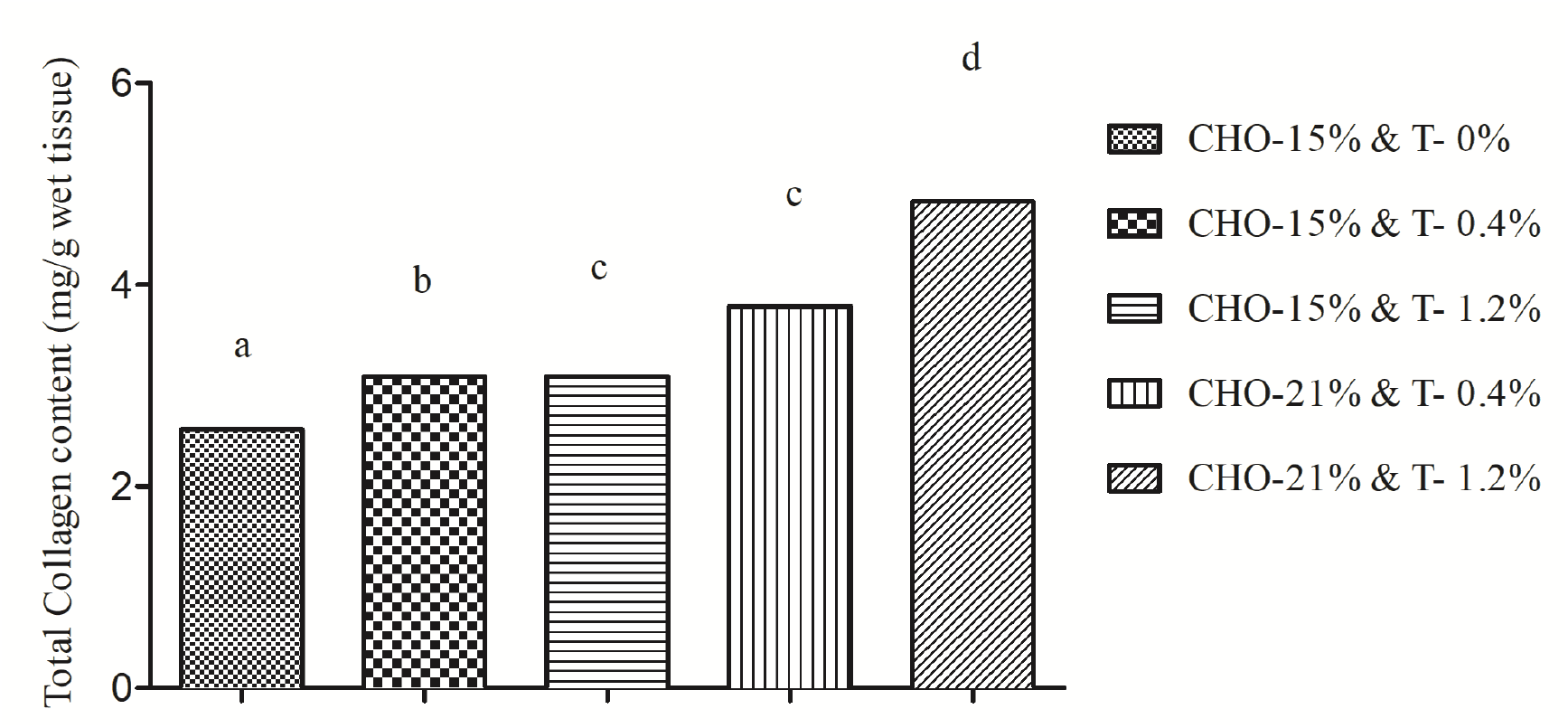
Total collagen content in the muscle of turbot after the 70-day feeding trial. superscript letters indicate the significant difference among the groups at 0.05 (*P* < 0.05). If the effects were significant, ANOVA was followed by the Tukey test.

### 3.4 Amino acid contents

Different treatment groups are affected differently by the levels of amino acids in the turbot skeletal muscle (Table 3). Threonine, Serine, Glutamine, Leucine, Phenylalanine, Isoleucine, Lysine, Histidine, and Arginine were significantly high in the H1 group than that in other all groups (*P* < 0.05). Glycine, Alanine, Valine, Tyrosine, and Proline were significantly highest in the H2 group (*P* < 0.05). So, most of the dietary essential amino acid levels have significantly increased by the 0.4% taurine and 21% carbohydrate levels fed turbot (*P* < 0.05). Even though muscle Cysteine was not significantly affected by either taurine or carbohydrates supplementation, the liver Cysteine level was significantly increased with the increased taurine inclusion and also by the carbohydrate levels in the two-way ANOVA analysis (*P* < 0.05).

**Table 3.** Amino acid composition (% dry matter) in the muscle of turbot fed with different levels of dietary taurine.

| Amino Acid | C | L1 | L2 | H1 | H2 |
| --- | --- | --- | --- | --- | --- |
| Aspartate | 10.66 ± 0.04<sup>c</sup> | 10.46 ± 0.16<sup>c</sup> | 10.80 ± 0.04<sup>c</sup> | 8.29 ± 0.03<sup>a</sup> | 9.33 ± 0.05<sup>b</sup> |
| Threonine | 4.84 ± 0.05<sup>a</sup> | 4.78 ± 0.04<sup>a</sup> | 4.86 ± 0.06<sup>a</sup> | 6.75 ± 0.02<sup>b</sup> | 4.97 ± 0.04<sup>a</sup> |
| Serine | 4.31 ± 0.06<sup>ab</sup> | 4.19 ± 0.07<sup>ab</sup> | 4.36 ± 0.08<sup>b</sup> | 5.27 ± 0.03<sup>c</sup> | 4.07 ± 0.04<sup>a</sup> |
| Glutamine | 17.18 ± 0.05<sup>c</sup> | 16.50 ± 0.13<sup>b</sup> | 17.5 ± 0.02<sup>c</sup> | 20.20 ± 0.24<sup>d</sup> | 15.15 ± 0.05<sup>a</sup> |
| Glycine | 3.98 ± 0.71<sup>a</sup> | 4.41 ± 0.07<sup>a</sup> | 4.86 ± 0.03<sup>ab</sup> | 6.18 ± 0.03<sup>bc</sup> | 7.16 ± 0.09<sup>c</sup> |
| Alanine | 5.96 ± 0.04<sup>a</sup> | 5.83 ± 0.03<sup>a</sup> | 6.13 ± 0.04<sup>a</sup> | 7.26 ± 0.04<sup>b</sup> | 8.17 ± 0.15<sup>c</sup> |
| Cysteine | 0.49 ± 0.06<sup>a</sup> | 0.54 ± 0.05<sup>a</sup> | 0.38 ± 0.05<sup>a</sup> | 0.55 ± 0.02<sup>a</sup> | 0.53 ± 0.01<sup>a</sup> |
| Valine | 4.85 ± 0.07<sup>ab</sup> | 4.80 ± 0.06<sup>a</sup> | 5.03 ± 0.03<sup>bc</sup> | 5.45 ± 0.02<sup>d</sup> | 5.19 ± 0.01<sup>c</sup> |
| Methionine | 2.41 ± 0.05<sup>a</sup> | 2.68 ± 0.01<sup>b</sup> | 2.73 ± 0.07<sup>b</sup> | 2.34 ± 0.02<sup>a</sup> | 2.71 ± 0.07<sup>b</sup> |
| Isoleucine | 4.79 ± 0.05<sup>a</sup> | 4.72 ± 0.03<sup>a</sup> | 4.99 ± 0.01<sup>b</sup> | 5.12 ± 0.03<sup>b</sup> | 4.80 ± 0.04<sup>a</sup> |
| Leucine | 8.31 ± 0.05<sup>ab</sup> | 8.27 ± 0.08<sup>a</sup> | 8.49 ± 0.02<sup>b</sup> | 9.00 ± 0.03<sup>c</sup> | 8.27 ± 0.02<sup>ab</sup> |
| Tyrosine | 0.00 ± 0.00<sup>a</sup> | 0.01 ± 0.00<sup>a</sup> | 0.07 ± 0.04<sup>a</sup> | 0.00 ± 0.00<sup>a</sup> | 0.34 ± 0.05<sup>b</sup> |
| Phenylalanine | 4.76 ± 0.04<sup>a</sup> | 4.59 ± 0.05<sup>a</sup> | 4.77 ± 0.05<sup>a</sup> | 5.13 ± 0.03<sup>b</sup> | 5.21 ± 0.02<sup>b</sup> |
| Lysine | 7.47 ± 0.04<sup>b</sup> | 7.47 ± 0.04<sup>b</sup> | 7.55 ± 0.01<sup>b</sup> | 7.97 ± 0.01<sup>c</sup> | 6.68 ± 0.02<sup>a</sup> |
| Histidine | 1.73 ± 0.04<sup>ab</sup> | 1.66 ± 0.01<sup>a</sup> | 1.66 ± 0.03<sup>a</sup> | 1.82 ± 0.01<sup>b</sup> | 1.64 ± 0.02<sup>a</sup> |
| Arginine | 6.67 ± 0.06<sup>b</sup> | 6.40 ± 0.01<sup>a</sup> | 6.84 ± 0.01<sup>c</sup> | 7.30 ± 0.01<sup>d</sup> | 6.52 ± 0.04<sup>ab</sup> |
| Proline | 2.59 ± 0.03<sup>ab</sup> | 2.52 ± 0.02<sup>b</sup> | 2.84 ± 0.03<sup>c</sup> | 0.96 ± 0.04<sup>a</sup> | 4.72 ± 0.12<sup>d</sup> |

Notes: Data are expressed as mean ± SE. Mean values within the same row with different superscripts (a, b, c, d and e) are significantly different ( $P < 0.05$ ; Tukey's test).

### 3.5 Muscle texture

Adhesiveness, chewiness, cohesiveness, hardness, and springiness have shown the changes in textural properties of turbot muscle (Table 4). Hardness has significantly increased by all the treatment groups than that in the control group (*P* < 0.05). The highest value of hardness recorded in the H1 group which was supplemented with 0.4% taurine and 21% carbohydrates levels. There was significantly difference between 15% and 21% carbohydrate levels for the hardness of the turbot muscles which means the carbohydrate supplementation significantly increased the effect of taurine on muscle hardness (*P* < 0.05). But adhesiveness, chewiness, cohesiveness, and springiness were not significantly affected by any treatment group than that in the control group (*P* > 0.05). There was no significant interaction between carbohydrates and taurine in the two-way ANOVA analysis (*P* > 0.05).

**Table 4.** Muscle texture of turbot after the 70-day feeding trial.

| Muscle textural Properties | C | L1 | L2 | H1 | H2 |
| --- | --- | --- | --- | --- | --- |
| Adhesiveness (mJ) | 3.64 ± 0.04 | 3.90 ± 0.02 | 4.10 ± 0.32 | 3.96 ± 0.14 | 4.23 ± 0.90 |
| Hardness (g) | 90.47 ± 5.19<sup>a</sup> | 125.63 ± 2.67<sup>b</sup> | 138.23 ± 3.93<sup>b</sup> | 169.67 ± 8.93<sup>c</sup> | 146.23 ± 4.93<sup>bc</sup> |
| Cohesiveness | 0.40 ± 0.02 | 0.37 ± 0.02 | 0.40 ± 0.01 | 0.38 ± 0.02 | 0.42 ± 0.02 |
| Springiness (mm) | 0.54 ± 0.02 | 0.70 ± 0.03 | 0.62 ± 0.01 | 0.63 ± 0.06 | 0.64 ± 0.03 |
| Chewiness (mJ) | 43.30 ± 7.51 | 56.34 ± 1.82 | 46.25 ± 6.04 | 54.46 ± 2.85 | 47.75 ± 4.83 |

All data were expressed as mean ± SE. Mean values within the same row with different superscripts are significantly different ( $P < 0.05$ ; Tukey's test).

### 3.6 Digestive enzymes

Taurine and carbohydrates treated groups significantly affected the digestive enzyme activity more than that in the control group (*P* < 0.05) (Table 5). All the treated groups have significantly increased the amylase, lipase, and trypsin enzyme activities in the intestine of the turbot (*P* < 0.05). Amylase and lipase activities were significantly highest in the H2 group and it was not dependent on the carbohydrate levels (*P* < 0.05) and it was not dependent on the carbohydrate levels in two-way ANOVA analysis. There was no significant difference between the treatment groups for trypsin activity in the intestine (*P* > 0.05).

**Table 5.**
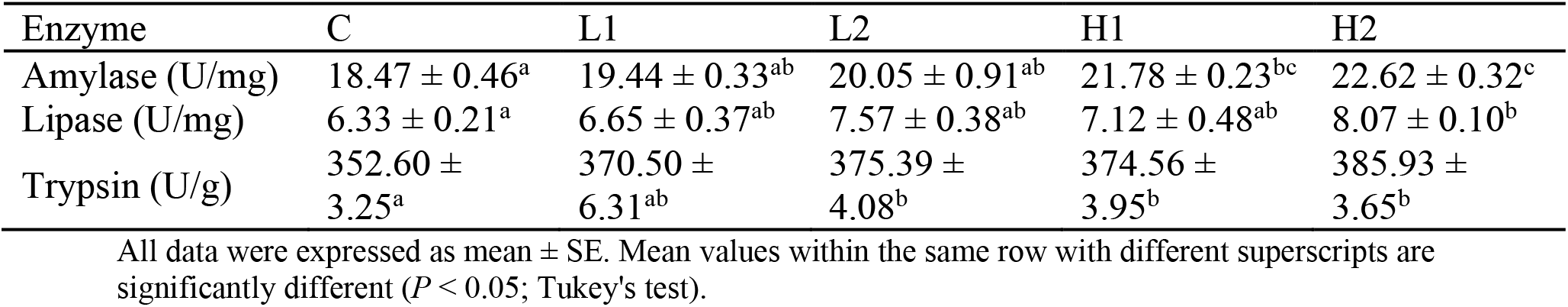
Effects of taurine and carbohydrate supplementation on digestive enzyme activities in the intestine of turbot after the 70-day feeding trial.

| Enzyme | C | L1 | L2 | H1 | H2 |
| --- | --- | --- | --- | --- | --- |
| Amylase (U/mg) | 18.47 ± 0.46<sup>a</sup> | 19.44 ± 0.33<sup>ab</sup> | 20.05 ± 0.91<sup>ab</sup> | 21.78 ± 0.23<sup>bc</sup> | 22.62 ± 0.32<sup>c</sup> |
| Lipase (U/mg) | 6.33 ± 0.21<sup>a</sup> | 6.65 ± 0.37<sup>ab</sup> | 7.57 ± 0.38<sup>ab</sup> | 7.12 ± 0.48<sup>ab</sup> | 8.07 ± 0.10<sup>b</sup> |
| Trypsin (U/g) | 352.60 ± 3.25<sup>a</sup> | 370.50 ± 6.31<sup>ab</sup> | 375.39 ± 4.08<sup>b</sup> | 374.56 ± 3.95<sup>b</sup> | 385.93 ± 3.65<sup>b</sup> |

All data were expressed as mean ± SE. Mean values within the same row with different superscripts are significantly different ( $P < 0.05$ ; Tukey's test).

## 4 Discussion

Taurine is a sulfur amino acid that is mainly available in animal tissues. It has mainly concentrated in skeletal muscle, heart, brain, retina, blood cells, large intestine, and plasma of the mammalian tissues. Dietary taurine has increased the muscle growth, muscle amino acid contents, cell volume homeostasis, nutrition metabolism, enzyme activities, muscle composition, antioxidative capacity, immunity, survival, liver bile acid production, feed efficiency, broodstock and larval performance of the fish (El-Sayed, 2013). Even though taurine is a non-essential nutrient, its inclusion in diets is often recommended because of its auxiliary actions such as membrane protection, antioxidation, and detoxification in mammals and fish (Zhang et al., 2018; Abdel-Tawwab and Monier, 2017), osmoregulation in invertebrates and higher animals (Schaffer et al., 2000), carrier of lipid-soluble vitamins in mammals (Petrosian and Haroutounian, 2000) and enhancement of bile salt production in fish (Kim et al., 2015). Even though taurine is synthesized in the liver from methionine and cysteine, taurine biosynthesis ability varies according to the fish species. A taurine deficiency can lead to poor production performances and physiological abnormalities in different life stages of the fish (Shen et al., 2017).

Hyperplastic muscle fibers represent the new muscle fiber generation or muscle growth. Hypertrophy muscle fibers represent the size increase of existing muscle fibers. Hyperplastic muscle fibers are responsible for high muscle density and both hyperplastic and hypertrophy muscle fibers are responsible for muscle textural properties. Hyperplastic growth has key roles at the juvenile stage of the fish. After that, the hypertrophic cellular growth takes place for further muscle growth of the fish muscle by enlarging the volume of the existing muscle fibers (Sampath et al., 2020b; Johnston, 1999; Rowlerson and Veggetti, 2001). So, the current study has agreed on the positive effect of taurine for optimal muscle growth by increasing the hypertrophy muscle fiber generation of the juvenile stage of the fish. Furthermore, hyperplastic muscle fiber generation was stimulated by the high carbohydrate levels. Moreover, increased taurine and carbohydrate levels have increased the muscle’s total collagen by increasing alkaline-soluble hydroxyproline content and alkaline-insoluble hydroxyproline content. So, total collagen content is also facilitated muscle growth and muscle textural properties other than hyperplastic muscle fiber development of the turbot.

Individual amino acid supplementation has engaged with the ionic regulation of certain amino acids including glycine, taurine, arginine, proline, and alanine in shrimp and fish (Sampath et al., 2020b; El-Sayed, 2013). Dietary taurine affects the amino acid level of the fish. Taurine has increased the growth performance and feed utilization of Nile tilapia (*Oreochromis nilotictus*) larvae up to 1% of taurine supplementation content with the soybean meal-based diet. Meanwhile, body protein and body amino acids were significantly increased by taurine (Al-Feky et al., 2016). And also, dietary taurine supplementation in plant-based diets fed Senegalese sole (*Solea senegalensis*) has significantly increased lipid amino acid retention (*Solea senegalensis*) (Richard et al., 2017). The same taurine effects on different levels of amino acid levels in turbot muscle were observed in the current study. It has shown a significant increase in the amount of Aspartate, Threonine, Serine, Glutamine, Leucine, Phenylalanine, Isoleucine, Lysine, Histidine, and Arginine in turbot-fed 1.2% taurine and 15% carbohydrate levels. Intestinal amylase activity was significantly increased with dietary taurine supplementation with a low fish meal diet fed juvenile black carp (*Mylopharyngodon piceus*) (Zhang et al., 2018). And also, taurine has significantly decreased the catabolic enzyme activity of glucogenesis to facilitate the optimum carbohydrate metabolism in fish (Bañuelos-Vargas et al., 2014). Taurine has significantly increased the growth performance by increasing the effects of digestive enzymatic activities in the intestine of turbot (*Scophthalmus maximus*) Intestinal amylase, mRNA levels of liver glucose-6-phosphate dehydrogenase (G6PD), glucokinase, pyruvate kinase, phosphofructokinase, and glycogen synthase (GS) were significantly increased with 1.2% taurine and 21% carbohydrate levels supplementation of turbot (*Scophthalmus maximus L*.). (Zhang et al., 2019; Liu et al., 2017). The current study also confirmed the significantly increased intestinal amylase in 21% carbohydrate levels with 1.2% taurine. And also, there was no significant difference between the treatment groups for intestinal trypsin activity (*P* > 0.05). It confirmed the same results in Zhang et al. (2019) by suggesting the ability of taurine on intestinal enzyme activities and the stimulatory effect of carbohydrates on the metabolic function of taurine. In the case of lipase activity, taurine can emulsify lipids, to increase fat-soluble vitamin absorption, and enhance the dietary lipids by increasing bile acid synthesis in the liver (Magalhães et al., 2019). Taurine has significant effects on lipid metabolism, lipid digestion and the lipid absorption of the yellowtail (*Seriola quinqueradiata*) (Nguyen et al., 2013). The current study also confirms the significant effects of taurine on lipase activity in 21% carbohydrate levels with 1.2% taurine-fed juvenile turbot (*Scophthalmus maximus L*.). So, dietary taurine and carbohydrate supplementation have positively affected glucose metabolism, enzyme activity, and growth performance of turbot (*Scophthalmus maximus*). And also, taurine effects have been simulated by the combined effects of carbohydrates with taurine diets (Zhang et al., 2019).

## 5 Conclusion

The present study showed that taurine 1.2% with carbohydrates 21% treatment (H2) fed turbot significantly increased the hyperplastic muscle fiber generation, muscle fibers density, alkaline-soluble hydroxyproline content, alkaline-insoluble hydroxyproline content, total hydroxyproline, the total collagen, Glycine, Alanine, Valine, Tyrosine, and Proline. And also, the amylase, lipase, and trypsin enzyme activities in the intestine were significantly increased by taurine 1.2% with carbohydrates 21% treatment. Muscle quality fiber density was increased by increasing the hyperplastic muscle fiber generation. Total collagen content was increased by alkaline-soluble hydroxyproline content and alkaline-insoluble hydroxyproline of the muscle. Muscle hardness was significantly highest with 0.4% taurine and 21% carbohydrate levels. Hyperplastic muscle fiber generation was stimulated by the high carbohydrate levels. Finally, the muscle growth and quality of the muscle were increased by the optimum combination of hyperplastic muscle fiber, collagen content, amino acid, and intestinal digestive enzyme activities.

## Compliance and ethical statement

### Conflict of interest

The authors have declared that no conflict of interest.

### Ethics statement

The present study was designed according to guidelines of the Use of Experimental Animals of the Ocean University of China.

## Notes

### Competing Interest Statement

The authors have declared no competing interest.

